# Placental polyamines regulate acetyl-coA and histone acetylation in a sex-specific manner

**DOI:** 10.1101/2021.06.09.447192

**Authors:** Irving LMH Aye, Sungsam Gong, Giulia Avellino, Roberta Barbagallo, Francesca Gaccioli, Benjamin J Jenkins, Albert Koulman, Andrew J Murray, D Stephen Charnock-Jones, Gordon CS Smith

## Abstract

Fetal sex differences play an important role in the pathophysiology of several placenta-related pregnancy complications. We previously reported that the maternal circulating level of a polyamine metabolite was altered in a fetal sex-specific manner, and was associated with pre-eclampsia and fetal growth restriction. Here we show that placental polyamine metabolism is altered in these disorders and that polyamines influence widespread changes in gene expression by regulating the availability of acetyl-CoA which is necessary for histone acetylation. Sex differences in polyamine metabolism are associated with escape from X chromosome inactivation of the gene encoding the enzyme spermine synthase in female placentas, as evidenced by biallelic expression of the gene in female trophoblasts. Polyamine depletion in primary human trophoblasts impairs glycolysis and mitochondrial metabolism resulting in decreased availability of acetyl-CoA and global histone hypoacetylation, in a sex-dependent manner. Chromatin-immunoprecipitation sequencing and RNA–sequencing identifies downregulation of progesterone biosynthetic pathways as a key target and polyamine depletion reduced progesterone release in male trophoblasts. Collectively, these findings suggest that polyamines regulate placental endocrine function through metabolic regulation of gene expression, and that sex differences in polyamine metabolism due to XCI escape may buffer the effects of placental dysfunction in pregnancy disorders.

## Introduction

Maternal and perinatal deaths account for 6-7% of all deaths globally (W.H.O., 2008). Placental dysfunction is implicated in the etiology of many of the associated conditions, including preeclampsia and fetal growth restriction. Fetal sex influences both the incidence and outcome of pregnancies affected by these disorders. As a fetal organ, the placenta also exhibits sex-differences in its function. Although the underlying causes of placental sex differences are still incompletely understood, sex hormones and genetic factors are likely to play major roles.

The genetic and epigenetic factors that contribute to placental sexual dimorphism largely center on the X chromosome. The X chromosome can contribute to sex-differences in gene expression by means of escape from X chromosome inactivation (XCI). During XCI, one X chromosome is selected (randomly in humans) for transcriptional silencing. However, as many as 23% of X-linked genes have been reported to escape XCI in humans (depending on tissue type) (Tukiainen et al., 2017), and display biallelic expression. We previously purported that XCI contributes to the female-bias in placental expression of X-linked genes (Gong et al., 2018a). However, little is known about the functional consequences of the genes escaping XCI, especially as it relates to placental function and its relationship with pregnancy disorders.

Our previous study showed fetal sex-dependent associations in the polyamine metabolite N1,N12-diacetylspermine (DAS) in the serum of pregnant women with pregnancy disorders preeclampsia and fetal growth restriction (Gong et al., 2018a). Polyamines are ubiquitous polycations that are essential for cell growth, differentiation and survival (Pegg, 2016). At the molecular level, polyamines regulate gene expression through wide-ranging mechanisms including direct binding to nucleic acids, translation initiation, and interacting with transcription factors and epigenetic processes (Madeo et al., 2018).

In mammalian cells, polyamines consist of putrescine, spermidine and spermine. Putrescine is generated from ornithine by ornithine decarboxylase (ODC) in the rate-limiting step for polyamine biosynthesis. Putrescine is converted to spermidine and subsequently to spermine via their respective enzymes spermidine- and spermine-synthase (SRM and SMS). Polyamine homeostasis is achieved through tight regulation of their synthesis as well as through catabolism by enzymes including spermidine/spermine acetyltransferase (SSAT) which acetylates spermidine and spermine facilitating their cellular export.

Polyamine biosynthesis is upregulated during placentation (Lopez-Garcia et al., 2009), and knockout or pharmacological inhibition of ODC in rodents results in embryo lethality (Pendeville et al., 2001) and abnormal placental development (López-García et al., 2008). Moreover, human infants with loss-of-function mutations in spermine synthase (SMS), a condition known as Snyder Robinson Syndrome (SRS), are born small-for-gestational age (SGA) (Albert et al., 2015; Becerra-Solano et al., 2009). Despite these profound phenotypes, how impairments in polyamine metabolism cause these effects is still unclear.

In this study, we tested the hypothesis that SMS escapes XCI resulting in biallelic expression in female placentas resulting in sex-biased differences in polyamine metabolism. We further investigate the functional consequences of this sex-difference and show that polyamines control mitochondrial metabolism. Moreover, mitochondrial insufficiency induced by polyamine depletion reduced acetyl-CoA availability and histone acetylation resulting in altered expression of genes regulating progesterone synthesis and decreased progesterone secretion.

## RESULTS

### Sex differences in polyamine metabolism are associated with XCI escape of SMS

We previously showed that the female-bias in placental polyamine metabolism was associated with higher SMS expression in human placental tissues (Gong et al., 2018a). Moreover, we hypothesized that SMS escapes XCI based on indirect evidence showing female-biased expression and methylation patterns (Gong et al., 2018a). However, direct evidence of XCI escape requires demonstration of SMS expression from both copies of the X chromosome (i.e. active and inactive X). Therefore we investigated biallelic expression of SMS at the single cell level in trophoblast to account for random XCI. Because the syncytiotrophoblast layer, which makes up the majority of trophoblasts in the term placenta, is a multinucleated epithelium, isolation of single cells is not possible. Therefore we isolated single nuclei from placental tissues, and took advantage of heterozygous single nucleotide polymorphisms (hetSNPs) present within the mature mRNA of the SMS gene, enabling identification of the mRNAs encoded by the two copies of X chromosomes in females.

Using our placenta transcriptome data set (Gong et al., 2021), we identified four female placentas which contained a hetSNP in an exonic region of the SMS gene (position X:21940733 A/G). As an internal control, we identified an exonic hetSNP in TIMM17B (position X:48894188 G/A), an X-linked gene that was predicted to be X inactivated (i.e. does not show female-biased expression). The selected female placentas were heterozygous at both positions. Intact nuclei from each placenta were isolated, sorted into a single nucleus per well and the cDNA from each nucleus was obtained as illustrated in Figure 1A. To detect both alleles in each nucleus, duplex TaqMan SNP genotyping assays were performed whereby each allele-specific primer and probe set are labelled with different fluorescent dyes, one for each allele. To confirm that the predicted genotype inferred from the RNA-seq, the corresponding umbilical cord tissue was genotyped (Supplemental Figure 1A). Following multiplex qPCR analysis using allele-specific primers the individual nuclei were called as expressing the reference, alternate or both alleles. We confirmed that SMS was expressed from both alleles in single nuclei, with 4-40% of the individual nuclei exhibiting biallelic expression (Figure 1B and Supplemental Figure 1B). On the other hand, single nuclei expressed either the reference or the alternative TIMM17B alleles but never both in the same nucleus, confirming X inactivation of this gene (Figure 1B). These findings confirm that SMS escapes XCI resulting in biallelic expression in female placentas.

**Figure 1.**
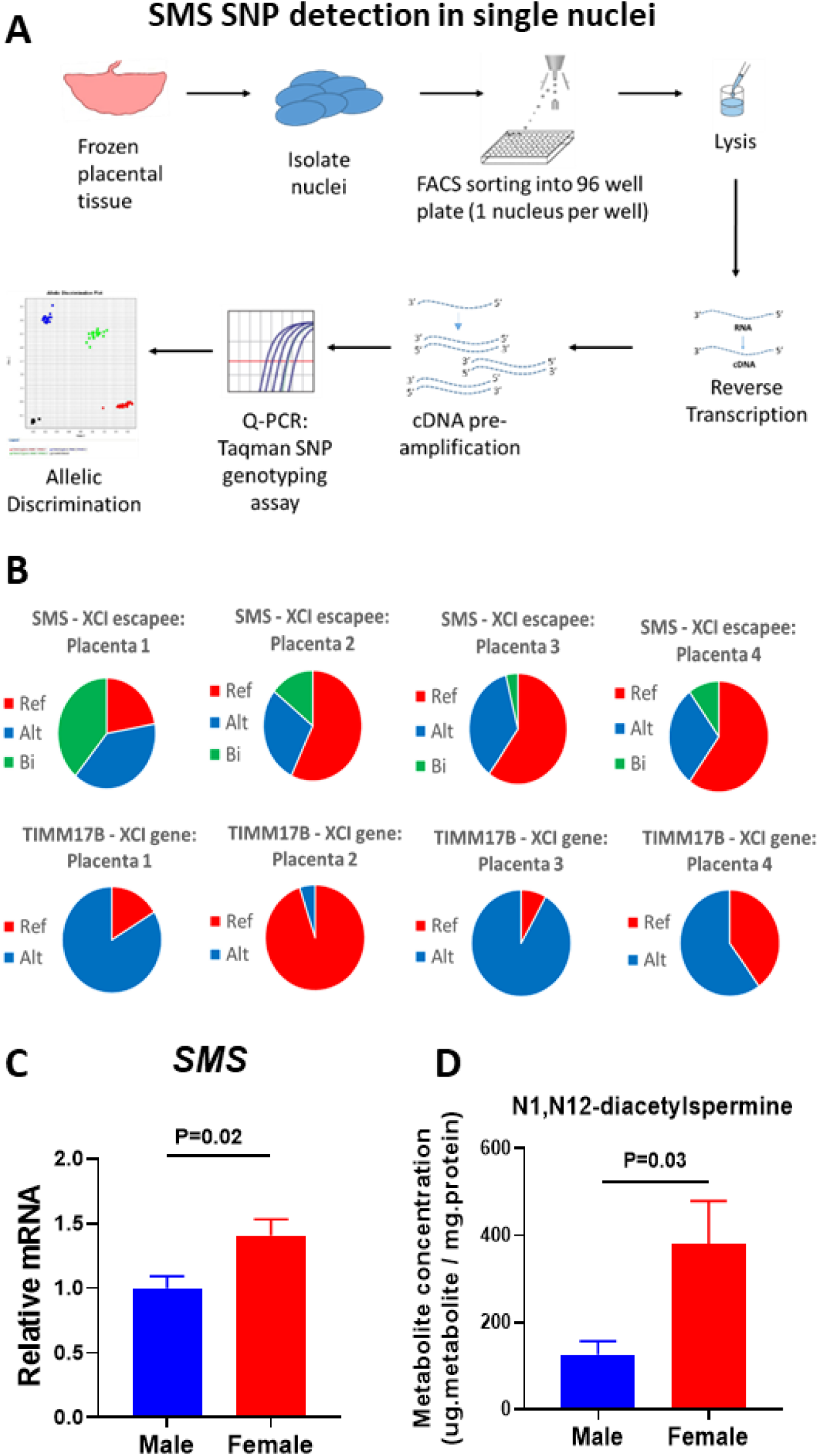
Sex differences in polyamine metabolism are associated with XCI escape of SMS. (**A**). Experimental workflow to assess XCI escape of SMS in single nuclei in female placental tissues. Nuclei were isolated from placental tissues and FACS sorted into 96 well plates with one nucleus per well. In the same well, nuclei were lysed, DNAse treated and mRNA reverse transcribed to cDNA, including a preamplification of the regions of interest. Resulting cDNA was then used for SNP typing using multiplexed taqman probes specific for both the reference and alternative alleles and allelic discrimination performed in each nuclei. (**B**) Monoallelic and biallelic expression of SMS (XCI escapee) and TIMM17B (XCI inactivated gene) SNPs in single placental nuclei. (**C**) SMS mRNA levels in PHTs. (**D**) Polyamine metabolite N1,N12-diacetylspermine concentrations in placental tissues. Bar graphs show mean±SEM, N=10 male PHTs and N=10 female PHTs.

We next explored the functional significance of female biallelic SMS expression by determining the mRNA level and activity of SMS using isolated primary human trophoblasts (PHTs). In female PHTs compared to males SMS mRNA was approximately 50% higher (Figure 1C) and the downstream polyamine metabolite DAS was 3-fold higher (Figure 1D). This demonstrates that the female biallelic SMS expression is associated with functional differences.

### Inhibition of polyamine metabolism profoundly alters the PHT transcriptome with enrichment of genes involved in mitochondrial metabolism

The specific role of polyamine metabolism on placental function was investigated by depleting polyamines from PHTs by inhibiting ornithine decarboxylase using difluoromethylornithine (DFMO). We previously showed that polyamine depletion with DFMO led to concentration-dependent reductions in PHT viability that were dependent on placental sex (Gong et al., 2018a). We chose the lowest concentration (5mM) of DFMO that mediated sex-specific responses for further analyses. DFMO treatment led to significant reductions in spermidine and spermine, although the reductions in spermine were greater in male PHTs (Supplemental Figure 2A).

To determine the effects of polyamine depletion on gene expression, we performed RNA-sequencing (RNA-seq) in PHTs following DFMO treatment. This analysis revealed profound changes in the PHT transcriptome of both sexes, with 4710 differentially expressed genes (DEGs) in male PHTs compared with 3558 DEGs in female PHTs (Supplemental Tables 1 and 2), with an overlap of 2689 DEGs between males and female PHTs (overlap P-value=1.7×;10^-69^). The RNA-seq results were confirmed by qPCR analysis of a subset of genes (corresponding to metabolic enzymes) in independent biological replicates (Supplemental Figure 2B and Supplemental Table 5).

To gain insights into the biological functions of the DEGs, gene set enrichment analysis (GSEA, (Subramanian et al., 2005)) was performed. Polyamine depletion in male PHTs resulted in significant enrichment (FDR q<0.05) of 9 pathways in the hallmark gene sets and 4 pathways in female PHTs (Supplemental Figure 3A) and 5 pathways in the KEGG gene sets (Supplemental Figure 3B) for male PHTs and 2 pathways for female PHTs. Genes associated with the tricarboxylic acid (TCA) cycle and Oxidative Phosphorylation (OXPHOS) were highly ranked in both the KEGG and Hallmarks gene sets (Supplemental Figure 3A and B), suggesting alterations in mitochondrial metabolism following polyamine metabolism.

To further investigate the link between polyamines and mitochondrial metabolism, we measured TCA cycle intermediates and polyamine metabolites in placental tissues from healthy pregnancies by liquid chromatography mass spectrometry (LC-MS) and performed correlation analyses. Significant positive correlations were observed between the polyamine end metabolites DAS (Figure 2A) and N1-acetylspermidine (NAS) (Figure 2B) and all 8 TCA cycle intermediates measured in both male and female placentas. These correlations were striking with values of Pearson’s r between 0.57 and 0.77 and adjusted P-values between 1.3×10^-10^ and 2.5×10^-21^.

**Figure 2.**
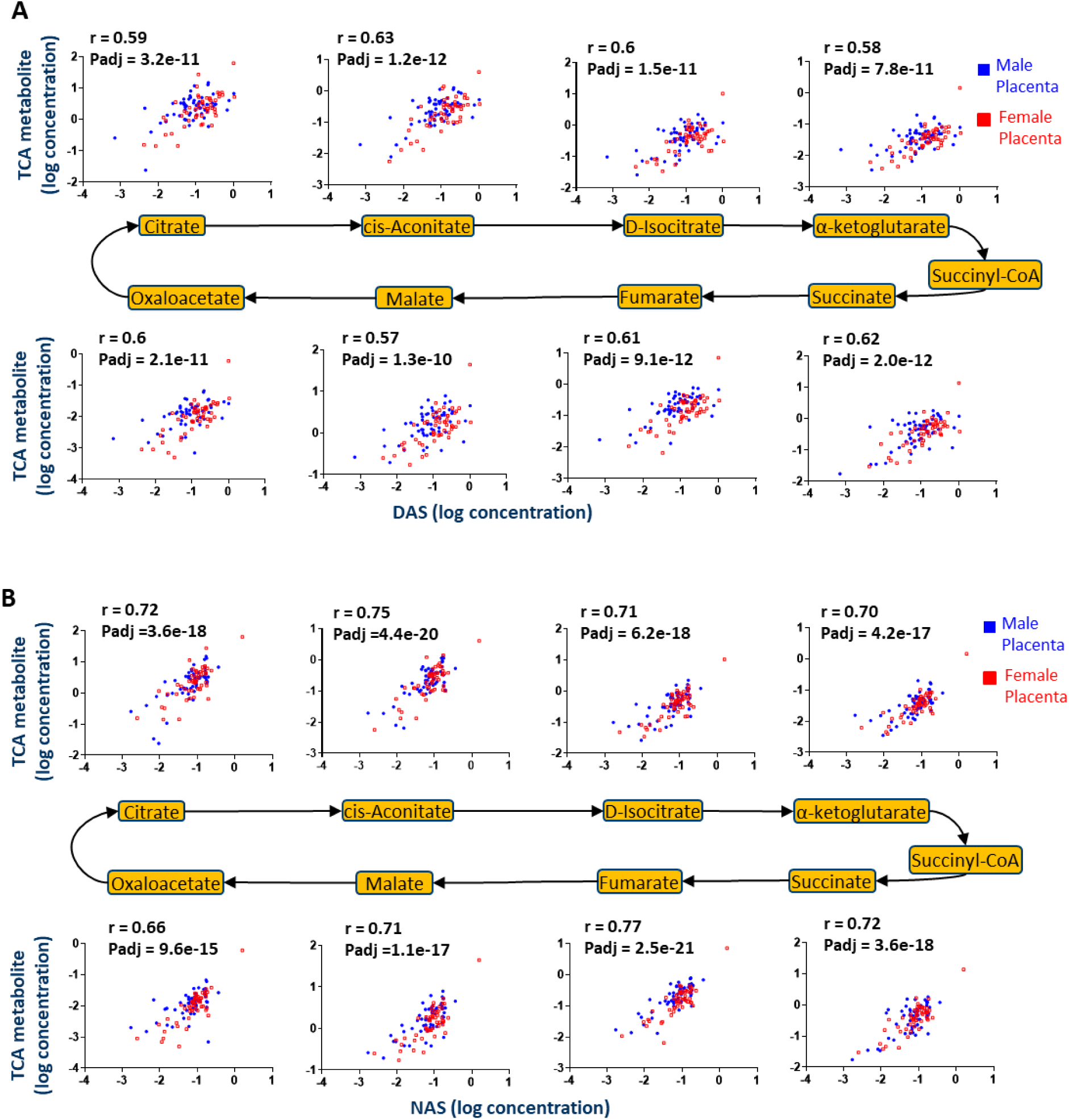
Correlations between TCA cycle intermediates and polyamine end metabolites. Scatter plots of TCA cycle intermediates versus (A) DAS or (B) NAS in human placental tissues. Metabolite concentrations were log transformed and Pearson’s correlation analyses performed. r = Pearson’s correlation coefficient. P-adj = P-values corrected for FDR by Benjamini-Hochberg method. DAS: N1,N12-diacetylspermine; NAS: N1-acetylspermidine. N=59 male placentas and N=51 female placentas.

### Polyamines regulate central energy metabolism

To clarify the causal relationship between polyamine metabolism and the TCA cycle, we measured the mRNAs encoding TCA cycle enzymes by qPCR and TCA cycle intermediates by LC-MS in PHTs depleted of polyamines (Figure 3A). Consistent with our RNA-seq results, DFMO significantly decreased the mRNA levels of 7 out of the 8 TCA cycle enzymes in male PHTs and 5 enzymes in female PHTs (Figure 3B). Similarly, polyamine depletion also led to significant reductions in TCA cycle intermediates in both male PHTs (6 out of 8 intermediates) and female PHTs (3 out of 8 intermediates) (Figure 3C).

**Figure 3.**
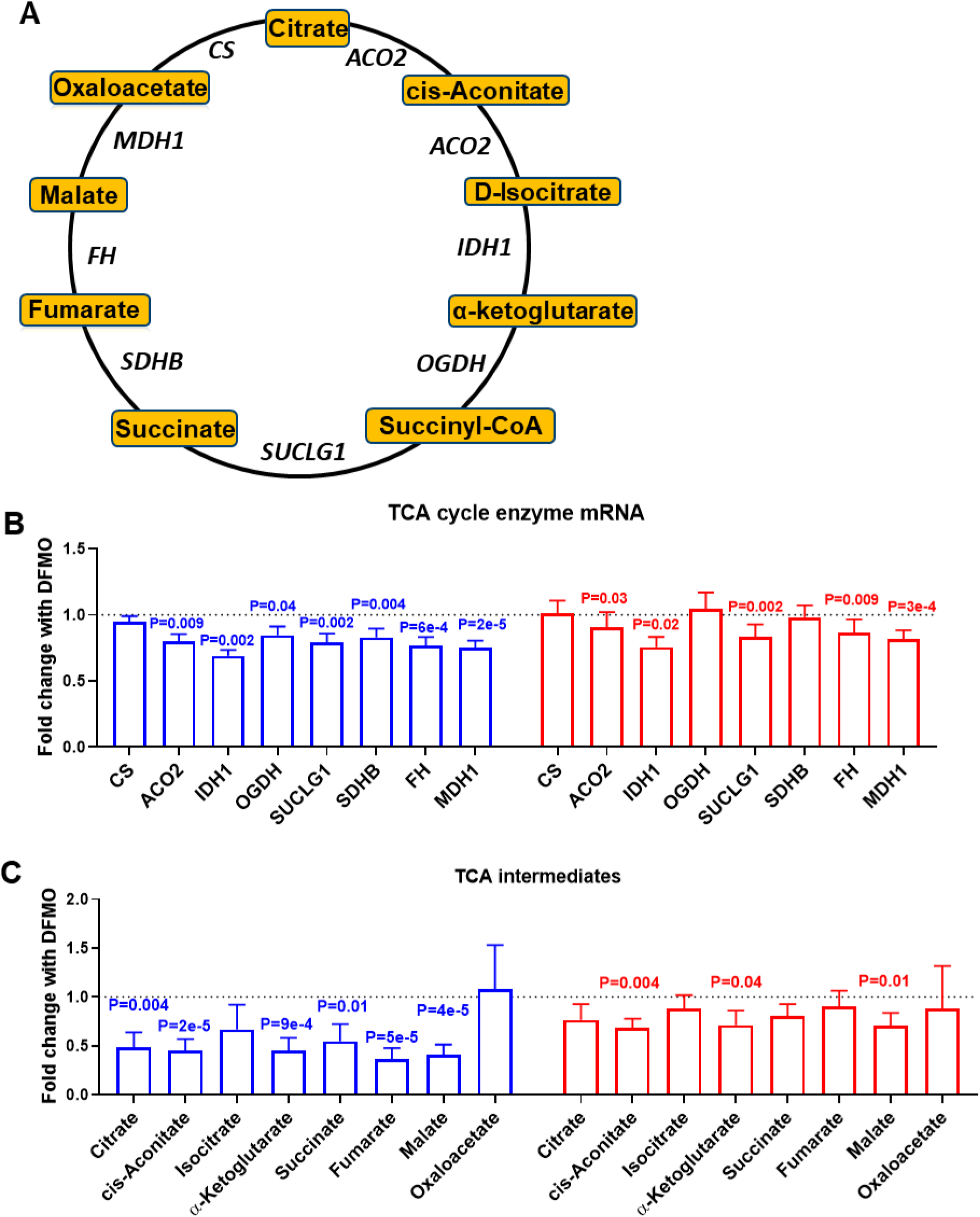
Polyamine metabolism is linked to TCA cycle metabolism. (A) Illustration of the TCA cycle indicating the metabolites and enzymes measured. (**B**) TCA cycle enzyme mRNA and (**C**) TCA cycle metabolites following DFMO-mediated polyamine depletion in PHTs. Bar graphs show mean±SEM, N=10 male PHTs and N=10 female PHTs.

We then determined whether the effects of polyamines extend towards functional changes in the two major cellular ATP-generating pathways, mitochondrial OXPHOS and glycolysis. Depletion of polyamines led to significant suppression in OXPHOS and basal glycolysis, although the sex-specific effects were dependent on the polyamine enzyme (Figure 4A-F). ODC inhibition with DFMO led to significant reductions in OXPHOS (Figure 4A) and glycolysis (Figure 4B) in male PHTs but not females (Supplemental Figure 4A).

**Figure 4.**
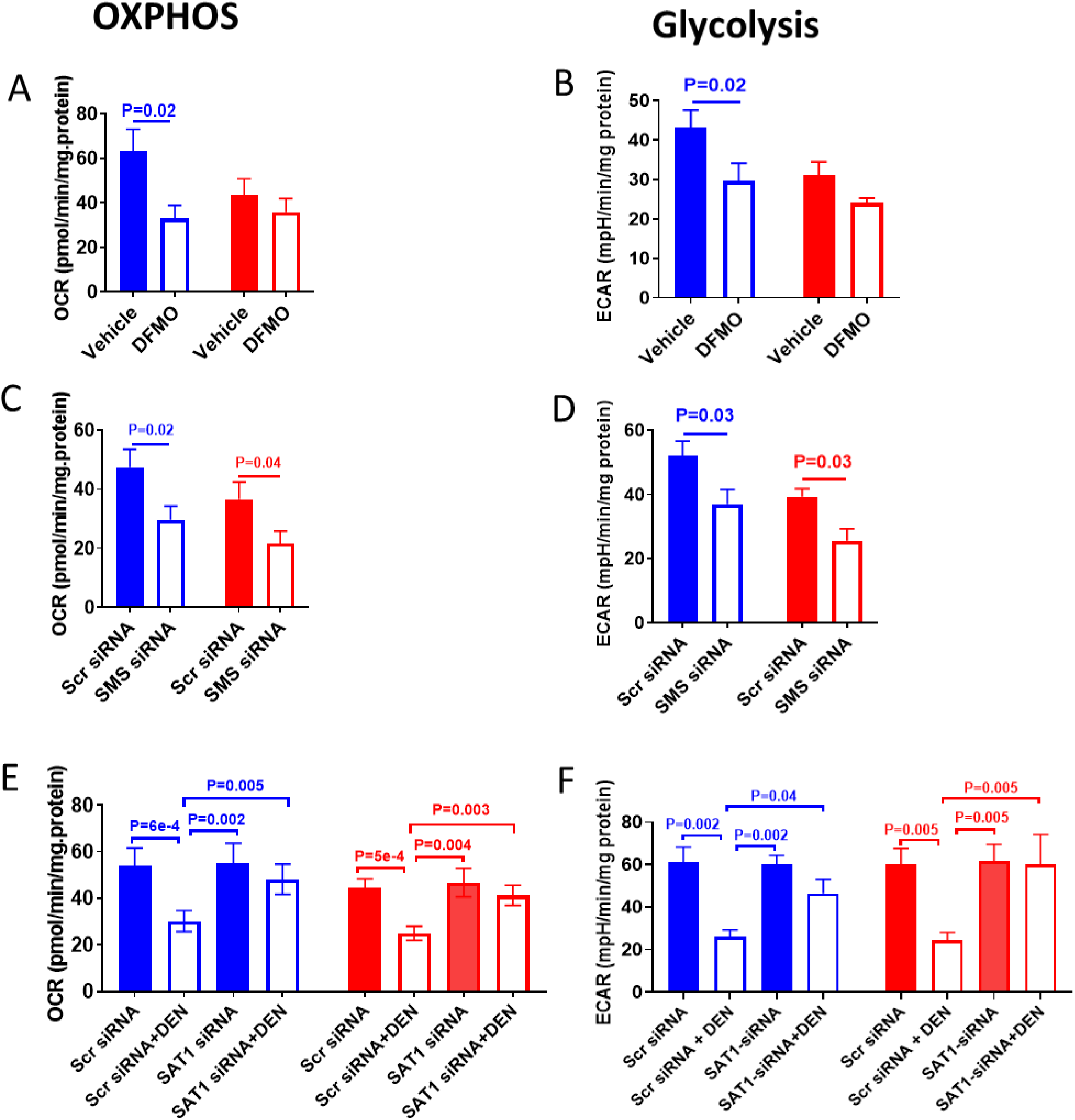
Suppression of cellular energy metabolism by polyamine depletion. Polyamine depletion by DFMO decreases (**A**) OXPHOS and (**B**) glycolysis in male PHTs. Inhibition of spermine synthesis by SMS silencing decreases (**C**) OXPHOS and (**D**) glycolysis in both male and female PHTs. Induction of SSAT-mediated polyamine catabolism by DENSPM decreases (**E**) OXPHOS and (**F**) glycolysis in both male and female PHTs. Bar graphs show mean±SEM, N=10 male PHTs and N=10 female PHTs. OCR, oxygen consumption rate; ECAR, extracellular acidification rate.

To confirm that the effects were due to polyamine depletion and not related to potential off-target effects of DFMO, we used additional approaches to deplete polyamines including SMS silencing (to inhibit spermine synthesis) and SSAT-induction with DENSPM (to activate polyamine catabolism (Fogel-Petrovic et al., 1997)). Supplemental Figures 4B and C show siRNA knockdown efficiencies. SMS-silencing reduced OXPHOS (Figure 4C) and glycolysis in both male and female PHTs (Figure 4D, and Supplemental Figure 4D). SSAT induction with DENSPM decreased OXPHOS (Figure 4E, and Supplemental Figure 4E) and basal glycolysis in both male and female PHTs (Figure 4F). These effects were reversed by SAT1 (gene encoding SSAT) siRNA confirming the involvement of SSAT induction in mediating DENSPM’s effects on energy metabolism (Figure 4E and 4F).

### Acetyl-CoA availability links polyamine metabolism to histone acetylation and gene expression

Previous studies have shown that cellular bioenergetic status is linked to transcriptional remodelling associated with histone acetylation through the availability of acetyl-CoA (Shi and Tu, 2015; Wellen et al., 2009). Histone acetyltransferases (HATs) utilise acetyl-CoA and this is their rate-limiting substrate. Thus, cellular acetyl-CoA levels profoundly influence HAT activity, and subsequently, histone acetylation. Given the profound changes in both gene expression and bioenergetics with polyamine depletion, we postulated that reductions in acetyl-CoA levels due to altered mitochondrial metabolism could lead to global hypoacetylation of histones in PHTs. Consistent with this hypothesis, DFMO significantly reduced acetyl-CoA levels in male but not female PHTs (Figure 5A). We then measured the lysine acetylation of several H3 histone marks by immunoblotting analyses following polyamine depletion. DFMO decreased H3K9 and H3K27 acetylation in male PHTs but not females, while H3K14Ac and H3K18Ac were not affected in either sex (Figure 5B). We focused on H3K27Ac because it is a robust mark of active promoters and distal enhancers that are tightly coupled to gene expression and transcription factor binding (Creyghton et al., 2010; Ernst et al., 2011), and previous studies had shown that alterations in this histone mark was associated with fetal growth restriction (Paauw et al., 2018). SSAT-mediated polyamine catabolism induced by DENSPM also reduced H3K27Ac abundance in both male and female PHTs, and this effect was reversed by silencing the SSAT-encoding gene SAT1 (Figure 5C).

**Figure 5.**
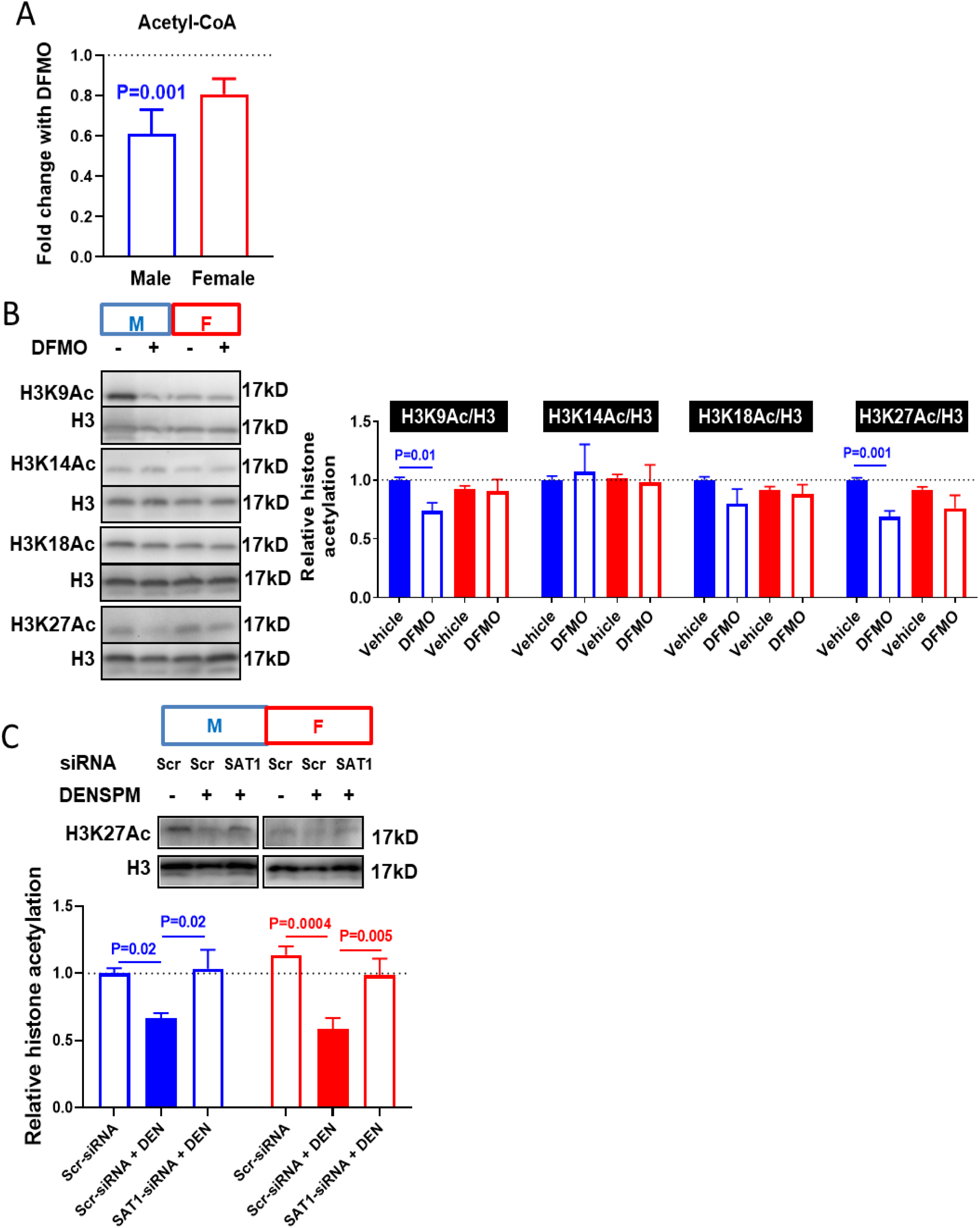
Polyamine depletion reduces acetyl-coA levels and decreases histone acetylation. DFMO decreases (**A**) acetyl-coA levels as measured by LC-MS and (**B**) histone acetylation as determined by immunoblotting, in male PHTs. (**C**) H3K27AC abundance is decreased by DENSPM (DEN) and reversed by SAT1 silencing. Bar graphs show mean±SEM, N=10 male PHTs and N=10 female PHTs.

Histone acetylation state is dependent on the balance between acetylation by HATs and deacetylation by histone deacetylases (HDACs). Therefore, the possibility also exists that polyamine depletion decreased histone acetylation by HDAC activation. To test this alternative hypothesis, we pre-treated PHTs with the global HDAC inhibitor (trichostatin A, TSA), followed by DFMO. We reasoned that if polyamine depletion by DFMO activated HDAC activity, then inhibition of HDACs would prevent the DFMO-mediated changes in histone acetylation (Supplemental Figure 5A). As shown in Supplemental Figure 5B, HDAC inhibition by TSA increased H3K27 acetylation that was partially reversed by DFMO treatment. This data suggests that the effects of polyamine depletion on histone acetylation are not mediated by HDAC activity, but rather by regulating acetyl-CoA availability.

### Genome-wide profiling of H3K27Ac demonstrates polyamine metabolism sensitive genes

We next identified which epigenomic loci were most sensitive to polyamine levels by performing H3K27Ac ChIP-seq analysis of DFMO-treated PHTs. We obtained a median of 15 million reads across all samples (ChIPs and Inputs) and found 71,231 consensus peak regions where the majority (>58%) were annotated within the promoter regions, i.e. within 2kb of TSS (Figure 6A and Supplementary Figure 6A). Unsupervised principal component analysis of all H3K27Ac peaks demonstrated clear separation with vehicle treated and DFMO treated male PHTs but a lack of separation by treatments in female PHTs (Figure 6B), suggesting a sex-biased effect.

**Figure 6.**
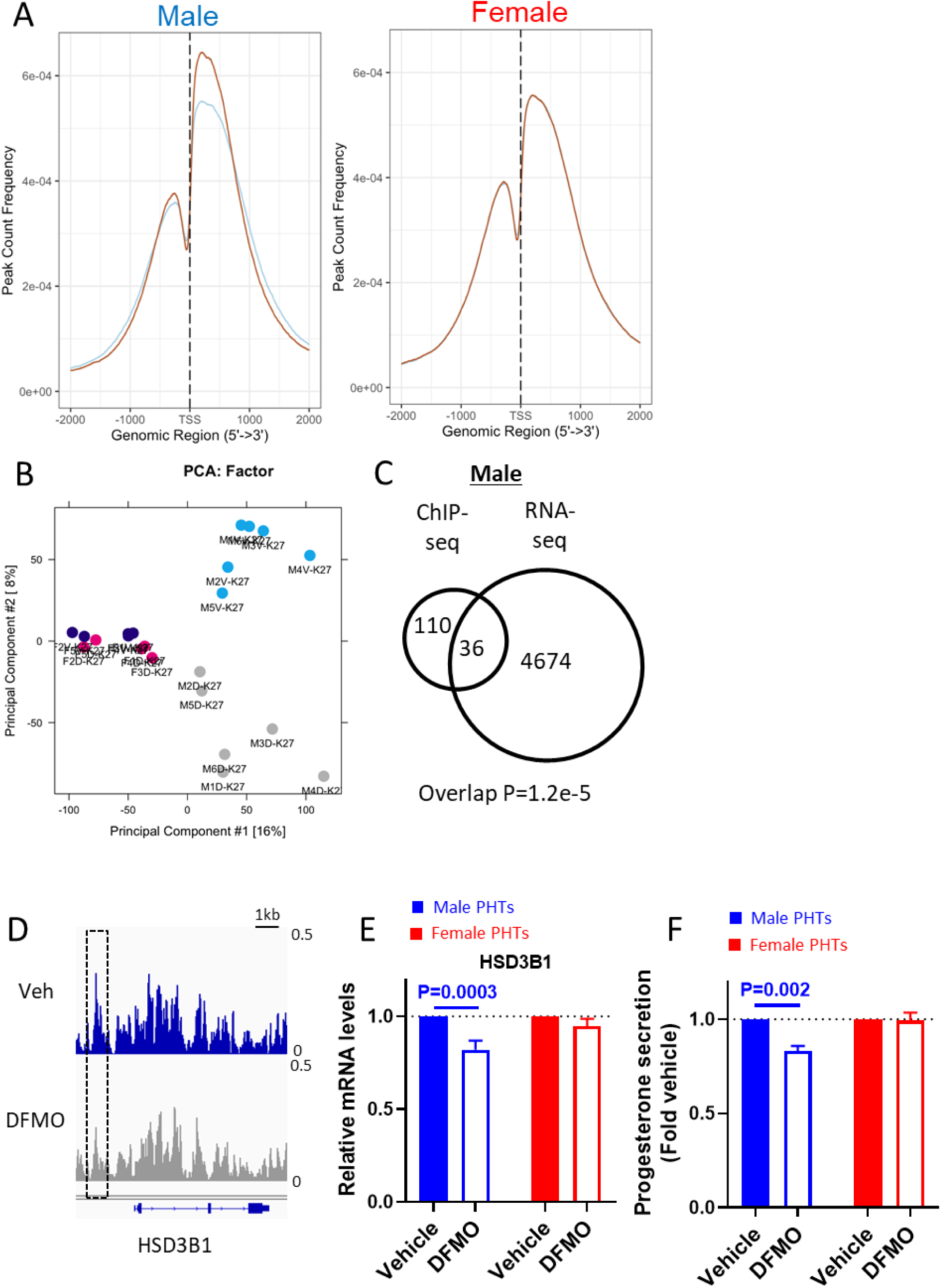
Polyamine depletion mediates global histone hypoacetylation and progesterone biosynthesis. (**A**) H3K27Ac binding near transcription start sites is decreased in male PHTs following DFMO treatment. Orange line represents vehicle, and blue line represents DFMO treated PHTs. (**B**) Unsupervised principal component analysis of H3K27Ac binding to genomic regions in vehicle and DFMO treated male and female PHTs. Blue dots show vehicle treated male PHTs, gray dots show DFMO treated male PHTs, violet dots show vehicle treated female PHTs, and pink dots show DFMO treated female PHTs. (**C**) Overlap of DBR annotated genes in H3K27Ac ChIP-seq analysis with DEG in RNA-seq analysis. (**D**) Custom views of differentially acetylated region in vehicle and DFMO treated male PHTs. Polyamine depletion decreased (**E**) HSD3B1 mRNA and (**F**) progesterone secretion in male PHTs. ChIP-seq experiments performed in N=6 male PHTs and N=5 female PHTs; Progesterone secretion measured in N=5 male PHTs and N=5 female PHTs. Male and female samples are colored blue and red respectively.

We identified differential bound regions (DBRs), separately for male and female PHTs, using DiffBind (v3.0.13) with so called ‘background’ normalisation method applied to edgeR and DESeq2 (see Methods for details). In male PHTs, we found 213 and 147 DBRs based on edgeR and DESeq2 respectively, representing 188 and 133 genes (Supplemental Tables 3 and 4). No DBRs were identified in female PHTs by edgeR or DESeq2, confirming the sex-biased effect of DFMO. Importantly, 90% of the male DBRs were associated with decreased acetylation, consistent with the global decreases in H3K27Ac levels examined by immunoblotting.

We then determined if these histone acetylation changes were associated with alterations in gene expression. We restricted our analyses to DBRs identified by both edgeR and DESeq2, thus resulting in 115 common DBRs. By integrating with RNA-seq data, 35 (∼30%) DBRs corresponded with altered mRNA levels (Figure 6C). We further excluded from our analysis, DBRs in intron or intergenic regions which resulted in 17 DBRs, of which 16 DBRs were downregulated by DFMO. From this restricted analysis, 3 of the downregulated DBRs and genes are involved in the regulation of steroid hormone biosynthesis. These include: INSIG1, which regulates cholesterol biosynthesis thereby providing the backbone for all steroid hormones; HSD3B1, an enzyme that catalyses pregnenolone to progesterone; and SLCO2B1 which transports conjugated steroid hormones (Supplemental Figure 6B). These DNA regions along with several other DBRs from our restricted analysis (VAC14, TGM2, ZDHHC1) were validated in independent biological replicates by ChIP-qPCR assays (Supplemental Figure 6B). We focused on HSD3B1 as this enzyme plays a major role in placental progesterone synthesis (Simard et al., 2005). Consistent with the reduced histone acetylation in the regulatory regions of HSD3B1 (Figure 6D), both HSD3B1 mRNA (Figure 6E) and progesterone release (Figure 6F) were significantly reduced by polyamine depletion in male PHTs but not female PHTs. Taken together, these findings suggest that global histone hypoacetylation mediated by polyamine depletion leads to physiological changes associated with altered endocrine activity.

### Dysregulation of placental polyamine metabolism by pregnancy complications

We lastly determined whether there was a relationship between placental polyamine metabolites and placental-related pregnancy complications using our previously described prospective cohort following a matched case-control study design (Gaccioli et al., 2017; Gong et al., 2021). NAS and DAS were measured in term placental tissues of women who delivered a baby with a customized birth weight centile less than 10^th^ percentile (small for gestational age, SGA) at term, and from women who developed pre-eclampsia (PE). These cases were matched one to one on the basis of maternal age, BMI, smoking status and mode of delivery to controls delivering a normally grown infant. Analysis of placental NAS and DAS concentrations from matched cases and controls indicated statistically significant changes in polyamine metabolites with SGA and PE. Both NAS and DAS were higher in placentas of both male and female SGA infants compared to their matching controls, whereas both NAS and DAS were higher only in male placentas from preeclamptic pregnancies (Figure 7).

**Figure 7.**
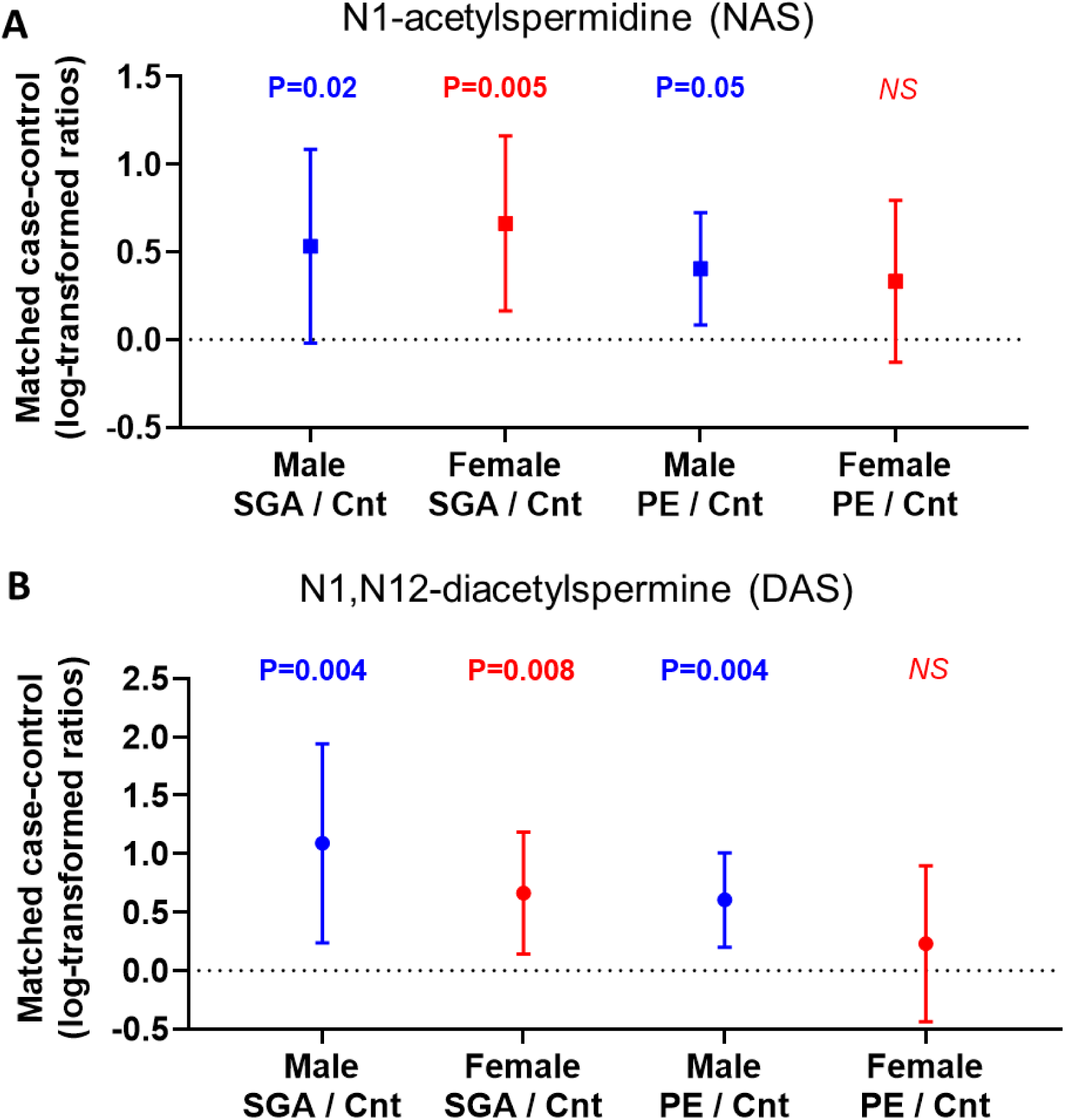
Placental polyamine metabolism is dysregulated in placental-related pregnancy complications. LC-MS analysis of DAS in placentas from pregnancies complicated with small-for-gestational age infants, preeclampsia and their matching controls. Scatter plot of differences of (**A**) N1-acetylspermidine (NAS) and (**B**) N1,N12-diacetylspermine (DAS) in case:control matched placental samples from pregnancies affected by SGA with a male fetus (N=16 pairs), SGA with a female fetus (N=18 pairs), PE with a male fetus (N=44 pairs) and PE with a female fetus (N=34 pairs). Mean ± 95% CI. Dashed line at 0 represents matching controls.

## Discussion

In this study, we provide several key insights into the mechanisms underlying sex-differences in placental function. To the best of our knowledge, this is the first report that an XCI escapee (SMS) contributes to the female-bias in placental function. Through a series of biochemical studies, transcriptomics and epigenomic profiling, our results suggest that the sex-bias in polyamine metabolism influences histone acetylation by regulating the availability of the acetyl donor, acetyl-CoA through its effects on mitochondrial metabolism. Moreover, using clinical samples from an optimally-phenotyped prospective cohort, we show that polyamine metabolism is dysregulated in the placentas of SGA and PE patients.

The higher DAS concentrations in preeclamptic placentas is consistent with our previous observations of increased DAS in the serum of preeclamptic women (Gong et al., 2018a). However, placentas complicated by SGA also show higher DAS whereas our previous report demonstrated reduced maternal serum DAS in pregnancies complicated by SGA. It is currently unclear why DAS levels in the placenta and maternal serum correlate in preeclamptic but not SGA pregnancies. We may speculate that the reduced placental size associated with SGA may have led to reduced maternal circulating levels of DAS. Alternatively, maternal renal failure in preeclampsia resulting in the inability to excrete DAS may have led to higher circulating levels of DAS, consistent with previous reports of elevated polyamine end-products in patients with chronic renal disease (Igarashi et al., 2006).

Our previous study predicted that 47 X-linked genes escape inactivation (Gong et al., 2018a), based on female-biased mRNA expression (i.e. higher levels in females compared to males) and DNA methylation (i.e. lack of promoter hypermethylation in females compared to males). These analyses identified SMS as a putative XCI escapee. However, investigating sex-biased expression and DNA methylation are proxies for XCI status and direct measurement of XCI can only be demonstrated at the single cell (or nucleus) level. By investigating XCI escape in single nuclei, we show biallelic SMS expression in female placentas that was present in up to 40% of cells, which is consistent with previous reports of heterogeneity in X-inactivation in human placentas (Phung et al.). Importantly, the biallelic SMS expression was associated with higher mRNA levels and greater concentrations of the polyamine end-metabolite DAS in female PHTs.

Polyamines are involved in many biological processes that are often varied and wide-ranging, and a complete mechanistic understanding of how polyamines exert their functions is presently lacking. Thus, we sought to clarify the role of polyamines and in particular, the functional significance of sex-differences in placental polyamine metabolism. We targeted several enzymes in the polyamine metabolic pathway using pharmacological and molecular approaches and found that polyamine depletion by DFMO was the only approach that produced sex-specific effects. As shown by the LC-MS analyses, DFMO decreased intracellular levels of spermidine and spermine (putrescine was not detectable) but the female-biased SMS expression was able to buffer some of the effect of polyamine biosynthesis inhibition and therefore female PHTs contained higher spermine concentrations compared to male PHTs following DFMO treatment.

Polyamine depletion led to large changes in gene expression, consistent with a role of polyamines in gene regulation (Childs et al., 2003). Pathway analyses indicated upregulation of inflammatory and stress signalling, and down-regulation of mitochondrial metabolism genes following polyamine depletion. The inflammatory and stress response is consistent with previous reports that polyamines prevent stress (Li et al., 2017; Rhee et al., 2007; Wei et al., 2016). However, the link to mitochondrial metabolism was intriguing as this has not been previously explored in detail. The strong correlations between acetylated polyamines and TCA intermediates suggests either that one of these metabolic pathways regulate the other, or that they are both under the control of a common upstream regulator. Polyamine depletion led to numerous changes in mitochondrial metabolism including decreases in TCA cycle intermediates and significant reductions in OXPHOS activity. The TCA cycle and OXPHOS are coupled since the reciprocal generation of NADH/FADH2 and NAD+/FAD respectively, fuels both processes (Martínez-Reyes and Chandel, 2020). Polyamines are mainly localised in the cytoplasm but rapid uptake of polyamines by the mitochondria has also been reported (Hoshino et al., 2005). It is therefore possible that cellular polyamine depletion also led to reductions in mitochondrial polyamines which altered mitochondrial function.

Alterations in mitochondrial metabolites can influence the epigenetic landscape by, for example, regulating the availability of substrates for histone modification enzymes. The Michaelis constant (Km) of HATs fall within the range of acetyl-CoA concentrations typically found in cells (Reid et al., 2017). Consequently, global levels of histone acetylation correlate with cellular acetyl-CoA abundance. In placental trophoblasts, acetyl-CoA is primarily generated in the mitochondria from pyruvate, the final product of glycolysis. Several findings from our study support a role for acetyl-CoA availability as the mechanism linking polyamine metabolism to histone acetylation. Glycolytic activity and acetyl-CoA levels were decreased in polyamine depleted PHTs, as were several histone acetylation marks, indicating global histone hypoacetylation. The histone acetylation effects were independent of HDAC activity. Lastly, ChIP-seq analysis of the most robustly decreased histone acetylation mark (H3K27Ac) showed predominantly decreased acetylation in numerous loci, consistent with a global hypoacetylation event. Induction of progressive mtDNA depletion which causes mitochondrial dysfunction resulted in decreased acetyl-CoA levels and global histone hypoacetylation (Lozoya et al., 2018). Overall, a picture emerges whereby polyamines have specific effects on the mitochondria that affect the histone acetylation landscape, but determining how this occurs requires further study.

Although histone acetylation explains some of the effects of polyamine depletion on gene regulation, given the large number of DEGs that were not accounted for by histone acetylation, additional factors are likely involved. Additional epigenetic effects such as histone methylation by α-ketoglutarate-dependent dioxygenases are plausible mechanisms linking polyamine-mediated mitochondrial dysfunction to histone modifications. Moreover, additional DNA-protein binding interactions also cannot be ruled out.

The locus-specific H3K27Ac and gene expression changes driven by polyamine depletion resulted in physiological outcomes as demonstrated by decreased acetylation and expression of HSD3B1, the enzyme critical for placental progesterone synthesis. Progesterone is essential for maintaining pregnancy and low levels of progesterone leads to miscarriage, and the placenta is the main source of progesterone after 10 weeks gestation (Johnson et al., 2018). Our findings are consistent with a previous study demonstrating H3K27 acetylation of HSD3B1 in PHTs (Kwak et al., 2019). Moreover, the reduced progesterone secretion by polyamine depletion in PHTs are also consistent with findings in mice showing a marked fall in progesterone following DFMO treatment (López-García et al., 2008). Therefore, our study establishes the mechanism linking polyamines and progesterone production by demonstrating that polyamines promote H3K27 acetylation and transcription of HSD3B1 leading to increased progesterone synthesis.

In summary, the results from this study support a model whereby SMS escapes XCI in female placentas which confers a protective effect against pregnancy complications. Mechanistically, our data suggest that polyamine mediated regulation of gene expression is determined, at least in part, by regulating acetyl-CoA availability which is necessary for histone acetylation. In future studies, it will be informative to explore the relationship between polyamines and central energy metabolism, and their link to epigenetic regulation.

Our findings may also extend towards broader evolutionary significance of XCI escapees. The Trivers-Willard hypothesis posited that under conditions of poor maternal resources, there is a survival advantage to invest resources in female offspring over male offspring (Trivers et al., 2011). Consistent with this hypothesis, clinical studies show that male fetuses that are growth restricted are more likely to be born stillborn than female fetuses (Smith, 2000), and therefore it may be less preferable to invest resources in males in a constrained pregnancy. Thus, the higher placental SMS expression in females due to XCI escape may be an evolutionary adaptation to buffer the effects of stress in female placentas in fetal growth restriction but not males.

## Methods

Please refer to supplemental table 6 for the complete list of key resources.

### Placental tissue collection

Placental tissues for primary human trophoblast cultures were collected from healthy women with normal term pregnancies and scheduled for delivery by elective Cesarean section. Participants were consented for research sample collection as part of the surgical procedure, with further permission for storage and transfer of materials to the biobank given under REC ID 07/MRE05/44. Analysis was performed as part of the Cambridge Blood and Stem Cell Biobank REC ID 18/EE/0199.

Metabolite analysis was performed in placentas from pregnancy complications and their matching controls using samples from the Pregnancy Outcome Prediction (POP) Study, previously described in detail (Gaccioli et al., 2017; Sovio et al., 2015). Ethical approval for the study was given by the Cambridgeshire 2 Research Ethics Committee (REC ID 07/H0308/163) and all participants provided written informed consent. Cases of preeclampsia (PE) were defined on the basis of the 2013 ACOG criteria (American College of Obstetricians and Task Force on Hypertension in Pregnancy, 2013) and cases of small for gestational age (SGA) had a customized birth weight <10th percentile (Freeman et al., 1995). Controls were defined as pregnancies resulting in a live born infant with a birth weight percentile in the normal range (20-80^th^ percentile (Freeman et al., 1995)) with no evidence of slowing in fetal growth trajectories, and with no evidence of hypertension at booking and during pregnancy, preeclampsia, Hemolysis/Elevated Liver enzymes/Low Platelet (HELLP) syndrome, gestational diabetes or diabetes mellitus type I or type II or other obstetric complications. A total of 222 unique placental samples were analyzed, among which there were 78 PE, 34 SGA, and 110 control samples.

### Primary human trophoblast (PHT) culture and targeting of polyamine metabolism

A total of 104 placentas were processed for primary human trophoblast (PHT) culture. PHTs were isolated by trypsin digestion and Percoll purification as previously described (Aye et al., 2010; Singh et al., 2012). Briefly, approximately 40g of villous tissue digested in trypsin (0.25%, Gibco) and DNAse I (325 Kunits/mg tissue, Sigma) and purified over a discontinuous 10–70% Percoll gradient centrifugation. Cells which migrated between 35–55% Percoll layers were collected and cultured in 1:1 mixture of Dulbecco’s modified Eagle’s medium (Sigma-Aldrich) and Ham’s F-12 nutrient mixture (Gibco) containing 10% fetal bovine serum, 50μg/ml gentamicin, 60μg/ml benzyl penicillin and 100μg/ml streptomycin (Sigma), and incubated in a 5% CO2 humidified atmosphere at 37°C. Following 18 hours of culture, attached cells were washed twice in warmed Dulbecco’s PBS and culture media was changed daily.

Pharmacological targeting of polyamine metabolism was achieved either by treatment with difluoromethylornithine (DFMO, 5mM) or N^1^,N^11^-Diethylnorspermine (DENSPM, 10µM) from 66-90h of culture. For siRNA-targeting, PHTs were transfected with 100nM of siRNAs targeting *SMS* or *SAT1* using Dharmafect2 transfection reagent at 18h of culture. All experimental analyses were performed at 90h of culture.

### Metabolic phenotyping with Seahorse Bioanalyzer

To measure mitochondrial oxygen consumption and extracellular acidification rates, a Seahorse XF96 analyzer was used for the Mito Stress Test according to the kit’s instructions. Briefly, PHTs were plated onto a Seahorse XFe96 microplate (Agilent) at 0.5 x 10^6^ cells/well following isolation. PHTs were maintained in culture as described above until 90h when OCR and ECAR assays were performed. 1hr before the assay, the medium was changed to XF base Media (Seahorse Bioscience, Agilent) supplemented with 17.5 mM glucose, 1 mM sodium pyruvate and 2mM L-glutamine (equivalent to the concentrations present in PHT culture media). Cell metabolic rates were measured using XF96 Extracellular Flux Analyzer (Seahorse Bioscience, Agilent). Oligomycin (2µM), carbonyl cyanide p-trifluoromethoxyphenylhydrazone (FCCP, 2µM) and rotenone + antimycin (0.5µM each) were sequentially added according to the experimental protocol (Hill et al., 2012). Each experimental condition was repeated in 8 wells (technical replicates) per plate for each placenta, and the mean of the technical replicates considered n=1.

### Metabolite measurements

PHTs grown in 10cm dishes at 10 million cells per dish, were used for the measurement of polyamine and TCA cycle metabolites. At the end of the experimental manipulations as described above, culture dishes were placed on ice, washed twice in D-PBS and cells pelleted. For polyamine analyses, cell pellets were spiked with spermidine-(butyl-d8) trihydrochloride (Sigma Aldrich, #709891) as an internal standard and extracted in pre-chilled (−20°C) acetonitrile. Following centrifugation to remove insoluble material, extracts were dried under nitrogen gas and reconstituted in 0.1% formic acid in H_2_O. For analysis of TCA intermediates, cell pellets were extracted in methanol and spiked with internal standards and dried down in a rotary evaporator. Extracts were then reconstituted in 10mM ammonium acetate in H_2_O. For placental biopsies, approximately 10mg of frozen tissue was homogenized on a Bioprep-24-1004 homogenizer prior to metabolite extraction as described above.

LC-MS was performed as previously described (Gong et al., 2018a). Briefly, chromatographic separation was achieved on a ACE Excel 2 C18-PFP (150mm * 2.1mm, 2 µm) LC-column with a Shimadzu UPLC system. The column was maintained at 55°C with a flow rate of 0.5ml/min. A binary mobile phase system was used with mobile phase A; 0.1% formic acid in water, and mobile phase B; 0.1% formic acid in acetonitrile. The gradient profile was as follows; at 0 minutes_0% mobile phase B, at 2.5 minutes_0% mobile phase B, at 5 minutes_100% mobile phase B, at 7.5 minutes_100% mobile phase B, at 7.6 minutes_0% mobile phase B, at 11 minutes_0% mobile phase B.

Mass spectrometry detection was performed on an Exactive orbitrap mass spectrometer operating in positive ion mode. Heated electrospray source was used, the sheath gas was set to 40 (arbitrary units), the aux gas set to 15 (arbitrary units) and the capillary temperature set to 250°C. The instrument was operated in full scan mode at 4 Hz from m/z 75–500 Da.

### Detection of biallelic SMS mRNA expression in single nucleus female placentas

We used the GATK pipeline (Freeman et al., 1995) to identify hetSNPs (heterozygous Single Nucleotide Polymorphisms) in exonic regions of SMS from the RNA-seq alignment data (i.e. BAM files) of our human placenta cohort (Gong et al., 2021). Briefly, the pipeline takes the following steps: 1) marking duplicate reads using ‘markDuplicate’ of Picard, 2) splitting reads that contain ‘N’s in their CIGAR string using ‘splitNRead’ of GATK (subsequent submodules from GATK hereafter), 3) realignment of reads around the indel using ‘IndelRealigner’, 4) recalibrating base quality using ‘BaseRecalibrator’, 5) calling the variants using ‘HaplotypeCaller’, and finally 6) counting reads by the reference and alternative bases of hetSNPs using ‘ASEReadCounter’. We also selected a X-linked gene that is subject to XCI (i.e. a negative control of SMS) based on the following conditions: 1) reasonable expression level (normalized read count > 100), 2) no significant change of transcript abundance by sex (P_adj_>0.4 and log2(Fold Change)<1.1), and 3) fulfilling aforementioned conditions for the placenta and 19 non-placental tissues from our previous study (Gong et al., 2018a). There were 8 such X-linked genes and we chose TIMM17B as a pre-designed Taqman SNP genotyping assay for this SNP was readily available. We identified the SMS SNP (position X:21940733A/G; dbSNP ID: rs34507903) and the TIMM17B SNP (position X:48894188G/A; dbSNP ID: rs1128363) in four female placentas.

Nuclei from the corresponding frozen placental tissues were isolated using EZ prep buffer (Sigma #NUC-101). Frozen placental biopsies (25mg) were homogenized using a 2 ml Kimble dounce tissue grinder and nuclei pellets resuspended in Nuclei Suspension Buffer (Clontech #2313A) before filtering through a 40μm cell strainer. The collected nuclei were then stained with 10μg/ml DAPI and diluted to 10,000 nuclei per ml in PBS (without Ca and Mg). Nuclei were sorted by FACS into single nucleus per well in a 96 well plate filled with nuclear lysis buffer. The single cell-to-Ct kit (ThermoFisher #4458327) was used to perform reverse transcription, cDNA preamplification using allele-specific primers and cDNA synthesis. Genotyping by qPCR was performed using pre-designed Taqman SNP genotyping assay for TIMM17B rs1128363 (assay ID:C 11611029_1_) and custom Taqman SNP genotyping assay for SMS rs34507903 and contain sequence-specific primers and VIC and FAM dye-labelled probes to allow for allelic discrimination.

### RNA-sequencing analysis

Total RNA was isolated from PHT lysates and genomic DNA removed using the RNeasy Plus Mini Kit (Qiagen). RNA quality was assessed using the RNA Nano kit on an Agilent 2100 Bioanalyzer (Agilent Technologies), and RNA samples with an RNA integrity number > 8 were deemed suitable for RNA-seq experiments. Total RNA libraries were prepared into an indexed library using TruSeq Stranded total RNA library prep kit (Illumina) according to the manufacturer’s instructions. cDNA libraries were validated using high sensitivity DNA chip on the Agilent 2100 Bioanalyzer and quantified using the KAPA Library Quantification Kit (Roche). Indexed libraries were pooled and sequenced with a 50-bp single-end protocol on a HiSeq4000 platform (Illumina) by the Cancer Research UK Cambridge Institute Genomics Core Facility.

The primer sequences and poor-quality bases were trimmed from the sequencing reads using cutadapt (Martin, 2011) after assessing the quality of reads using FastQC (v0.11.4). Transcript quantification was performed using Salmon (v0.9.1) against Ensembl 90 version (GRCh38) of transcript annotations. Relative transcript abundance was measured in RPKM (Read Per Kilobase of transcript per Million mapped reads) and differentially expressed gene analysis was performed using DESeq2 (v1.18.1) Bioconductor package (Love et al., 2014).

Pathway analysis was performed by gene set enrichment analysis (GSEA v.4.1.0) which uses predefined gene sets from the Molecular Signatures Database (MSigDB v7.4). Hallmark gene sets and C2 curated KEGG subset gene sets were used and list of ranked genes based on FPKM values. Enriched pathways with normalized enrichment scores <0.25 and false discovery rate <5% were considered significant.

### Chromatin Immunoprecipitation (ChIP) Sequencing and ChIP-qPCR

PHTs grown in 10cm dishes (10 million cells per dish) and treated with DFMO or vehicle, were cross-linked in 1% formaldehyde in PBS for 10min. Cross-linking was terminated by adding 125mM glycine for 5min, rinsed twice in cold D-PBS. Cells were then flash frozen as cell pellets and stored at −80°C. Chromatin shearing were performed using Diagenode Chromatin shearing kit for Histones (#C01020010) with modifications. Briefly, cell pellets were re-suspended in Lysis Buffer 1 and homogenized on a dounce homogenizer to release nuclei and pelleted by centrifugation. Dounce homogenization was repeated using Lysis Buffer 2 and released nuclei pelleted by centrifugation. Nuclei were then lysed in iS Buffer 1, and chromatin was fragmented to approximately 200-bp by sonication for 6 cycles of 30 seconds on and 30 seconds off, on a Diagenode Bioruptor Pico. A small aliquot (20µl) of sheared chromatin fragments were RNAseA and proteinase K digested, DNA purified on columns and the DNA size distribution examined using the Agilent 4200 TapeStation (Agilent Technologies). Chromatin concentrations were then quantified using a Qubit fluorometer prior to immunoprecipitation. The remaining sonicated chromatin (2µg) was immunoprecipitated with 10µg of H3K27Ac antibody (Abcam ab4729) or normal rabbit IgG in ChIP Buffer (CST) and incubated on a rotator at 4°C overnight, or 10% of chromatin set aside as the input fraction. ChIP-grade Protein G magnetic beads (CST) were then added to the mixture and incubated for a further 4h. The antibody-bound beads were washed, cross-linking was reversed and DNA fragments purified using QIAquick DNA purification kit (Qiagen).

ChIP-seq libraries were prepared on 5 ng of DNA from ChIP and input samples using the SMARTer ThruPlex DNA-seq kit along with DNA HT Dual Index Kit (both from Takara Bio) according the manufacturer’s instructions. Libraries were sequenced on a HiSeq4000 (Illumina) with single end reads of 50bp by the Cancer Research UK Cambridge Institute Genomics Core Facility. We obtained a median of 15M reads across 43 samples (22 ChIP and 21 input samples). The primer sequences and poor quality bases were trimmed from the sequencing reads using cutadapt (v2.7 with Python 3.6.8) [Martin and Wang 2011]. Trimmed reads were mapped to GRCh38 version human genome using bowtie (v1.1.2) and duplicated reads were discarded using sambamba (v0.7.1). On average, 72% of reads were uniquely mapped and they were used for subsequent analyses. The alignment files of input samples (i.e. DNA input without ChIP) were merged into four groups by taking the following two factors: 1) treatment types (DFMO or vehicle), and 2) the sex of PHTs. Then we identified peak regions of ChIP samples using MACS2 (v2.2.7.1) by matching the treatment type and the sex between ChIP and input samples to remove background signals. We used 148bp as the fragment size (--extsize) and 2.7×10^9^ as the effective genome size (--gsize). A peak is considered significant if P_adj_ <0.05. To increase specificity, peaks supported by at least two ChIP samples of the same treatment-sex group were used and they were finally merged across the four groups. A total of 71,231 consensus peaks were generated and they were annotated using ChIPseeker (v1.26). Finally, differential binding regions (DBRs) were detected, separately for male and female PHTs, using DiffBind (v3.0.13). We used the ‘background’ normalization by binning 15,000bp across the genome and applied the normalization factor to the full library size. We used a paired design with the following formular, which was subsequently used by DESeq2 and edgeR: ∼pair + treatment.

Results from the ChIP-seq analyses were validated in biological replicates using ChIP-qPCR. Following ChIP with H3K27Ac as described above, qPCR analysis was performed using primers targeted against DNA regions of the following genes: INSIG1, HSD3B1, SLCO2B1, TGM2, VAC14 and ZDHHC1, as these are either implicated in progesterone homeostasis and/or fulfil the criteria described in the results. Primers were designed based on enriched DNA regions identified by ChIP-seq analysis (see Supplemental Table 5 for primer sequences). The fold enrichment values were normalized to 10% input.

### Western blotting

PHTs were harvested in RIPA buffer (50mM Tris HCl, pH=7.4; 150mM NaCl; 0.1% SDS; 0.5% Na-deoxycholate and 1% Triton X100) containing protease inhibitors and phosphatase inhibitor cocktail 1 and 2 (1:100, Sigma). Protein lysates were separated on a 4-15% TGX gel (Biorad) under reducing and denaturing conditions, followed by transfer onto PVDF membranes. Amido Black (Sigma) staining was performed and membranes imaged for quantification of total protein normalization before blocking in 5% BSA in TBS-0.1%Tween (TBS-T). Membranes were subsequently incubated in primary antibodies at 4°C. Following washes in TBS-T, membranes were incubated with peroxidase-conjugated secondary antibodies and visualized by enhanced chemiluminescence using Clarity Western ECL substrate (Bio-Rad). Target protein levels were normalized against total protein (as determined by Amido Black staining).

### Quantification and statistical analyses

Data are presented scatter plots showing mean±95% confidence interval, scatter plots showing individual values or bar graphs showing means+SEM. Sample sizes (indicated in figures) always refers to biological replicates (i.e. PHTs or placentas from individual pregnancies). Statistical analyses were performed using GraphPad Prism9. Normal distribution was confirmed by Shapiro-Wilk test and log-transformed if required. Unless otherwise stated, two-way ANOVA with treatment and sex as independent factors was applied with Holm-Sidak’s multiple correction applied and adjusted P<0.05 was considered significant. Relationships between polyamine metabolites and TCA cycle intermediates, or between RNA-seq and qPCR analyses were determined using Pearson’s correlation coefficients and corrected for multiple comparisons by Benjamini-Hochberg method. Statistical methods for RNA-seq and ChIP-seq analyses are described in their respective Methods sections.

### Software and data availability

The software used in this study is listed in the key resources table (Supplemental Table 6). The RNA-Seq and ChIP-Seq datasets generated in this study are available in the European Nucleotide Archive (ENA www.ebi.ac.uk/ena) under the accession number PRJEB45391 (www.ebi.ac.uk/ena/data/view/PRJEB45391).

## Author Contributions

ILMHA, GA and RB conducted the experiments. SG performed bioinformatics analyses. BJJ and AK performed mass spectrometry experiments. ILMHA, DSCJ and GCSS designed the study. ILMHA, AJM, DSCJ and GCSS interpreted the data and wrote the manuscript. All authors approved the manuscript.

## Acknowledgements

We are grateful to: the participants in the POP study; Emma Cook, Katrina Holmes and Josephine Gill for technical assistance. Funding: The work was supported by a Centre for Trophoblast Research Next Generation Fellowship to Irving Aye and the NIHR Cambridge Comprehensive Biomedical Research Centre (BRC, United Kingdom; G1100221). Giulia Avellino is supported by a PhD Scholarship from the Centre for Trophoblast Research and the BRC. Giulia Avellino and Roberta Barbagallo were supported by the Erasmus Plus traineeship.

## Supplemental Figure Legends

**Supplemental Figure 1:**
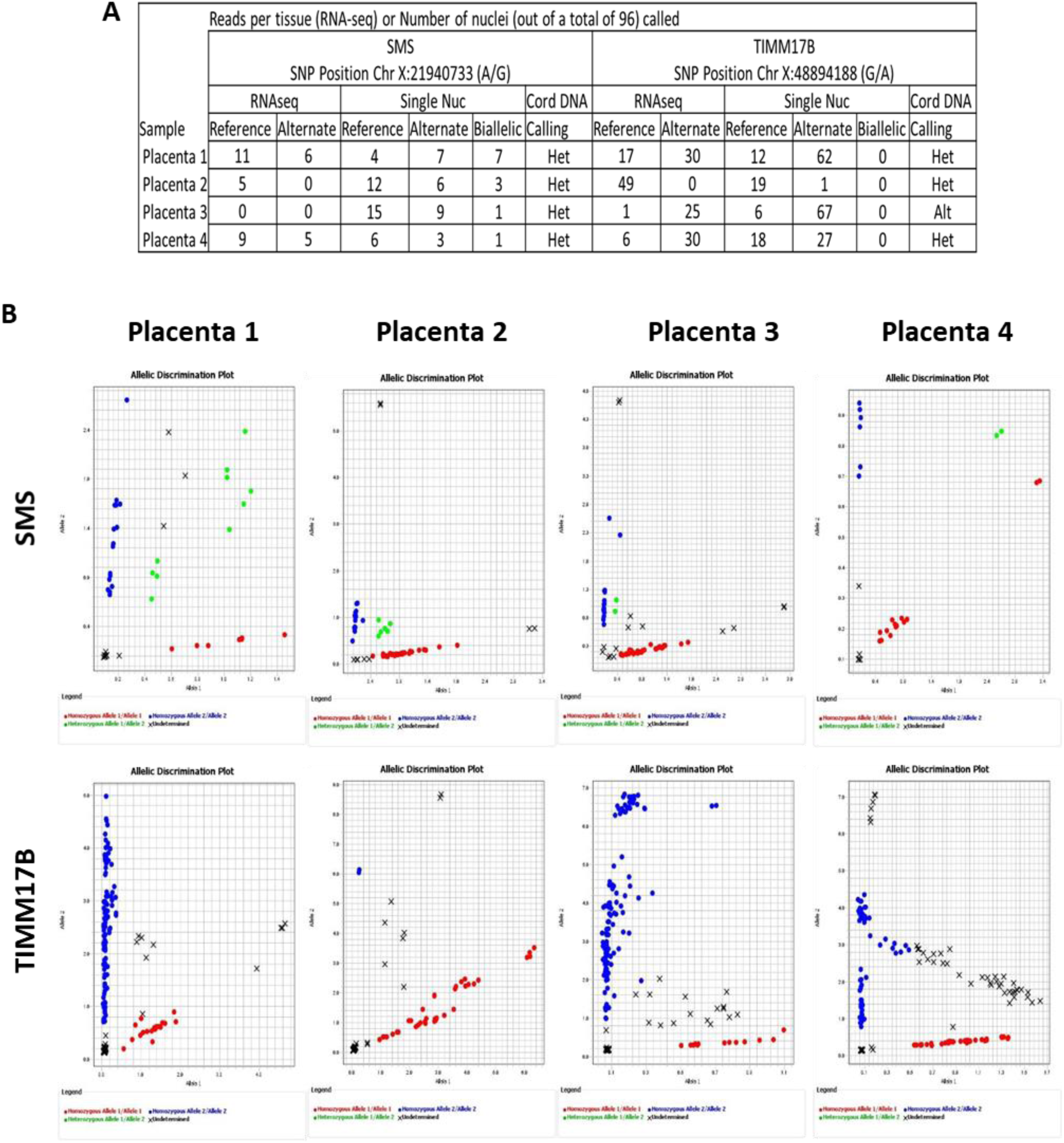
Allelic discrimination of SMS and TIMM17B in isolated single nuclei. (A) Allelic discrimination plots demonstrating reference, alternative or biallelic expression of SMS and TIMM17B in single nuclei. (**B**) Summary table of SNPs called in each placenta.

**Supplemental Figure 2:**
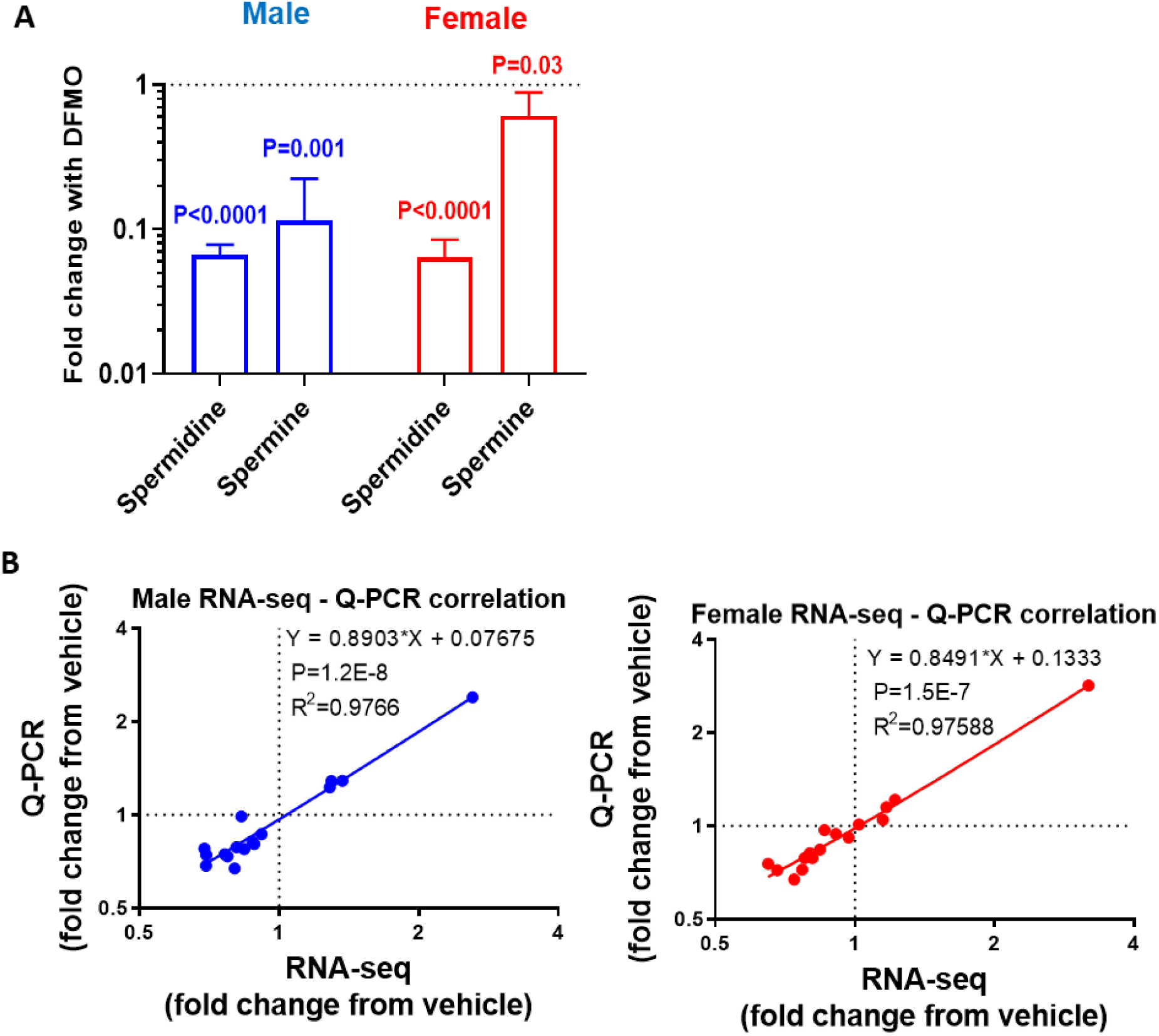
Sex-specific effects of DFMO on intracellular polyamine levels and RNA-seq validation. (**A**) Effect of DFMO on polyamine levels in PHTs. (**B**) Validation of RNA-seq by qPCR in independent biological replicates. Each data point represents a separate gene (mean fold change), N=10 male PHTs and N=10 female PHTs.

**Supplemental Figure 3:**
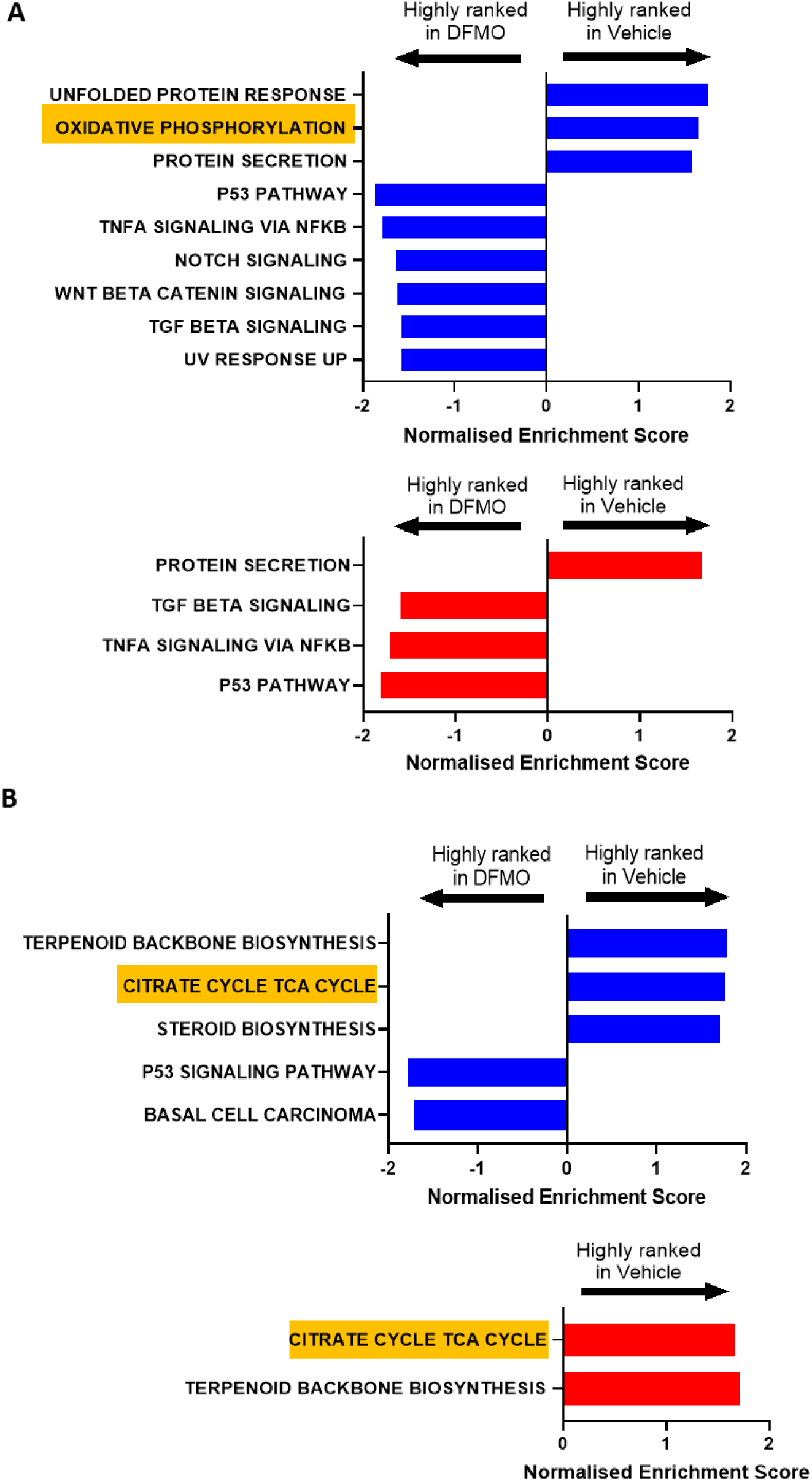
Gene set enrichment analysis of PHTs following polyamine depletion. Number of GSEA classes in which DEGs are overrepresented by DFMO treatment (FDR-adjusted *P-*value <0.05) in Hallmarks and KEGG pathways. (**A**) Normalised Enrichment Scores of HALLMARK pathways. (**B**) Normalised Enrichment Scores of KEGG pathways. DEGs highly ranked in DFMO indicate upregulation with DFMO and DEGs highly ranked in vehicle indicate downregulation with DFMO. Blue indicates pathways enriched in male PHTs and red indicates pathways enriched in female PHTs.

**Supplemental Figure 4.**
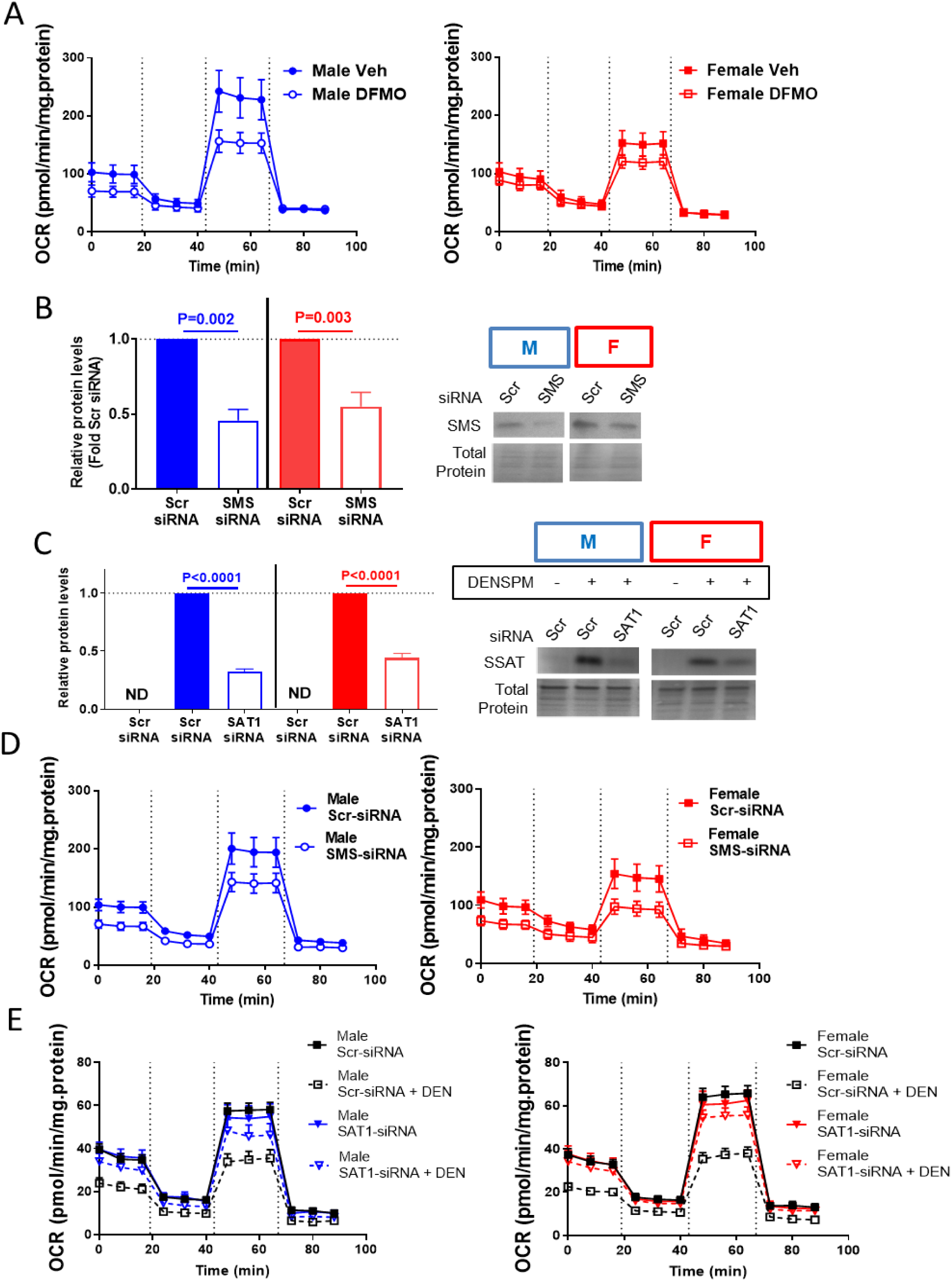
Oxygen consumption rate following targeting of polyamine metabolism. Oxygen consumption rate (OCR) in male and female PHTs following (**A**) DFMO treatment, (**B**) SMS silencing, (**C**) SSAT activation by DENSPM and reversal by SAT1 silencing. (**D**) SMS protein levels following siRNA silencing. (**E**) SSAT protein levels following DENSPM treatment with/without siRNA-silencing. Graphs show mean±SEM, N=10 male PHTs, N=10 female PHTs. ND, not detected.

**Supplemental Figure 5.**
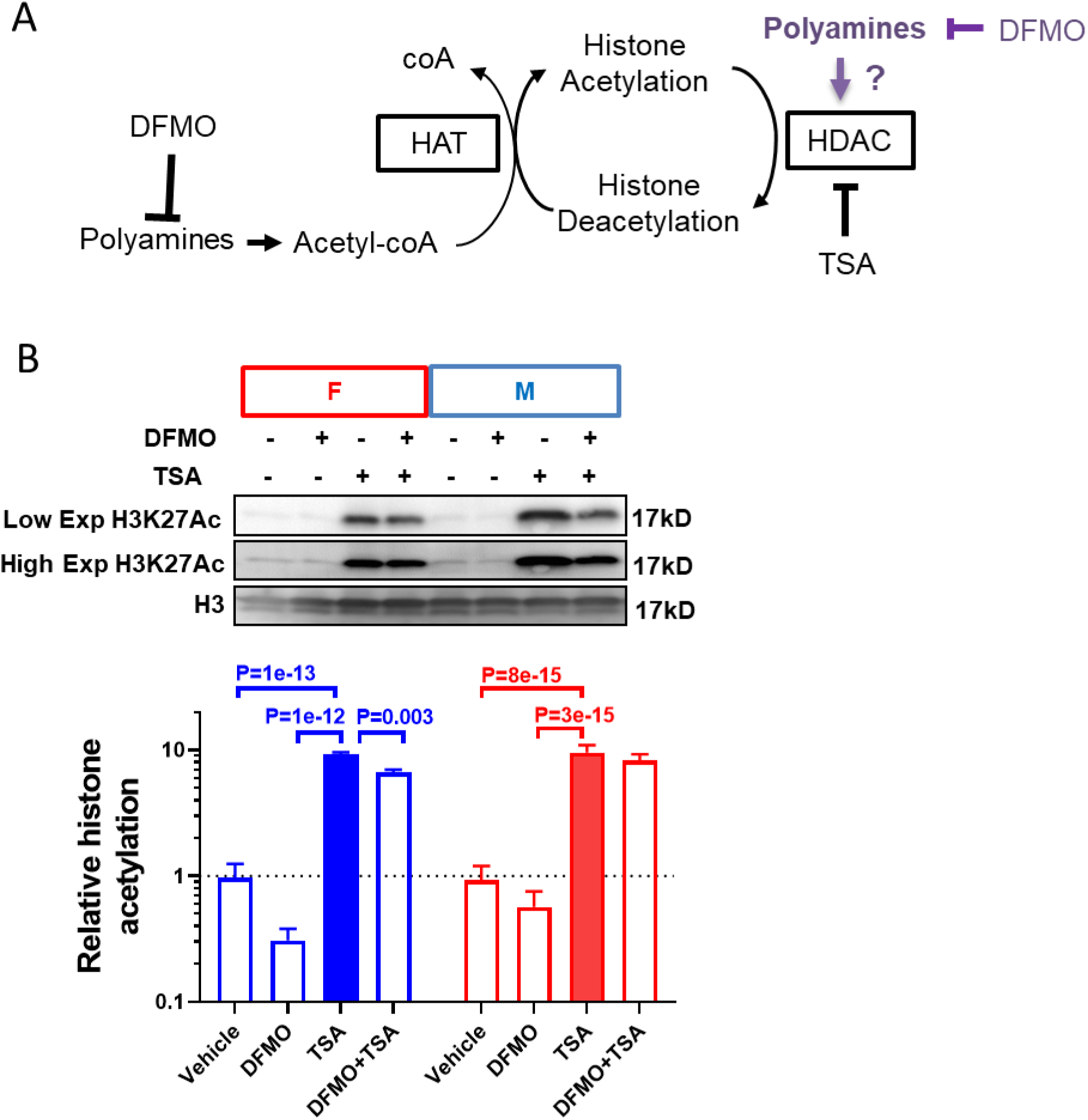
Histone deacetylase activity is not involved in polyamine’s effects on histone acetylation. (**A**) Model of the effect of polyamines on histone acetylation (shown in black) and alternative hypothesis (shown in purple). (**B**) Decrease in histone acetylation by DFMO is independent of HDAC inhibition by TSA. Bar graphs show mean±SEM, N=10 male PHTs and N=10 female PHTs.

**Supplemental Figure 6.**
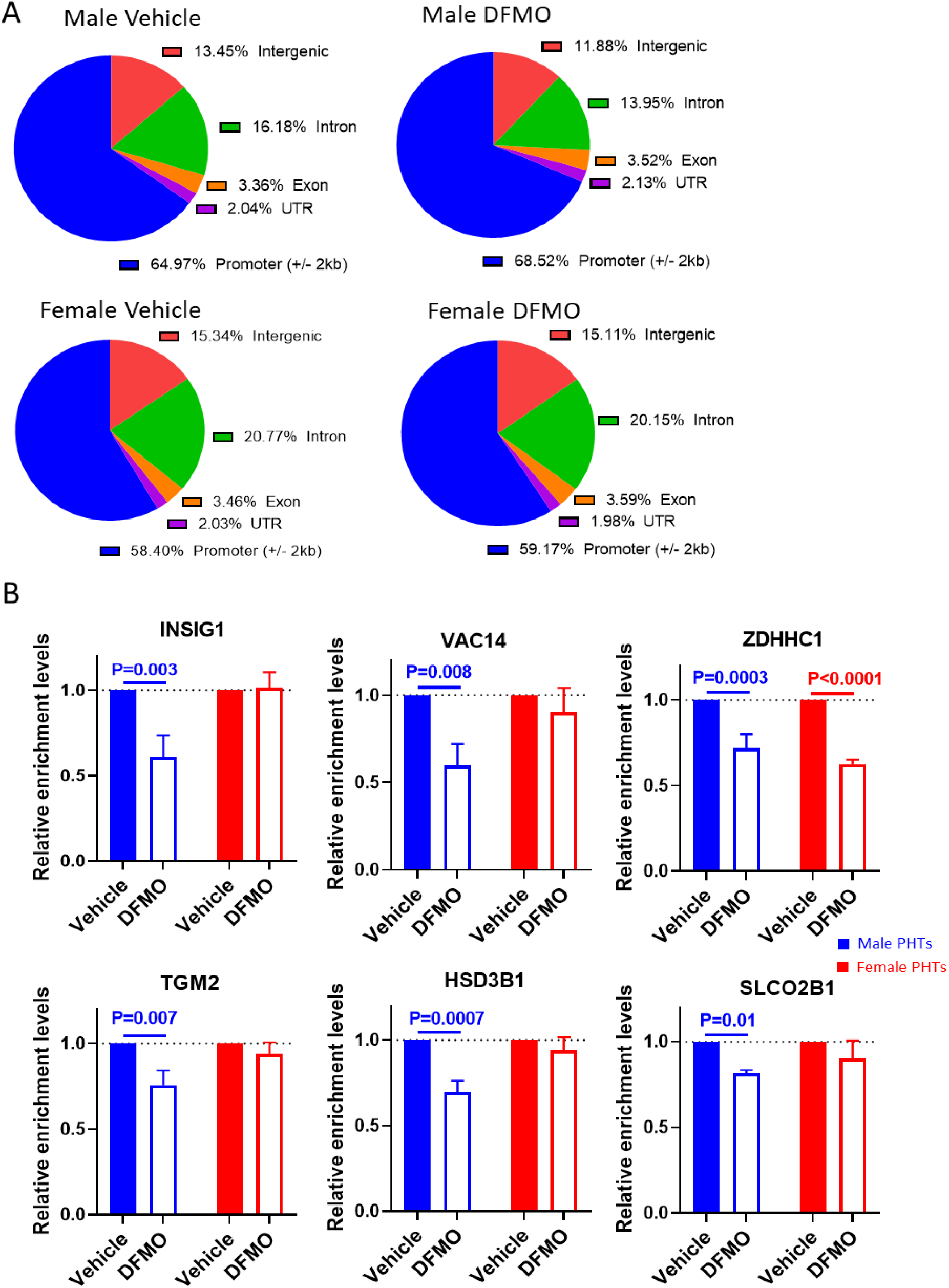
ChIP-seq and validation by ChIP-qPCR. (**A**) H3K27Ac occupancy in genomic regions of PHT. (**B**) Validation of ChIP-seq results by ChIP-qPCR in independent biological replicates. Bar graphs show mean±SEM, N=5 male PHTs and N=5 female PHTs.

## Notes

### Competing Interest Statement

The authors have declared no competing interest.

